# Monocyte-derived Dll4 is a novel contributor to persistent systemic inflammation in HIV patients

**DOI:** 10.1101/2023.04.18.537330

**Authors:** Shumin Wang, Meera Singh, Hongmei Yang, Craig N. Morrell, Laith Awad Mohamad, Jasmine Jiayuan Xu, Tiffany Nguyen, Sara Ture, Alicia Tyrell, Sanjay B. Maggirwar, Giovanni Schifitto, Jinjiang Pang

## Abstract

**Background:** In people living with HIV (PLWH) on combination antiretroviral therapy (cART), persistent systemic inflammation is a driving force for the progression of comorbidities, such as cardiovascular and cerebrovascular diseases. In this context, monocyte- and macrophage-related inflammation rather than T cell activation is a major cause of chronic inflammation. However, the underlying mechanism of how monocytes cause persistent systemic inflammation in PLWH is elusive.

**Methods and Results:** In vitro, we demonstrated that lipopolysaccharides (LPS) or tumor necrosis factor alpha (TNFα), induced a robust increase of Delta-like ligand 4 (Dll4) mRNA and protein expression in human monocytes and Dll4 secretion (extracellular Dll4, exDll4) from monocytes. Enhanced membrane-bound Dll4 (mDll4) expression in monocytes triggered Notch1 activation to promote pro-inflammatory factors expression. Dll4 silencing and inhibition of Nocth1 activation diminished the LPS or TNFα -induced inflammation. exDll4 releases in response to cytokines occurred in monocytes but not endothelial cells or T cells. In clinical specimens, we found that PLWH, both male and female, on cART, showed a significant increase in mDll4 expression, activation of Dll4-Notch1 signaling, and inflammatory markers in monocytes. Although there was no sex effect on mDII4 in PLWH, plasma exDll4 was significantly elevated in males but not females compared to HIV uninfected individuals. Furthermore, exDll4 plasma levels paralleled with monocytes mDll4 in male PLWH. Circulating exDll4 was also positively associated with pro-inflammatory monocytes phenotype and negatively associated with classic monocytes phenotype in male PLWH.

**Conclusion:** Pro-inflammatory stimuli increase Dll4 expression and Dll4-Notch1 signaling activation in monocytes and enhance monocyte proinflammatory phenotype, contributing to persistent systemic inflammation in male and female PLWH. Therefore, monocyte mDll4 could be a potential biomarker and therapeutic target of systemic inflammation. Plasma exDll4 may also play an additional role in systemic inflammation but primarily in men.

## Introduction

The World Health Organization (WHO) reported that approximately 38.4 million people live with HIV (PLWH) around the world in 2021. Although combination antiretroviral therapy (cART) has led to a substantial decline in the death rate of PLWH due to HIV infection, the death rate of non-communicable illnesses especially cardiovascular disease (CVD)^1^ has been rising. Compelling evidence demonstrates persistent systemic^2^ and local^3^ (such as coronary arteries) inflammation, a key mediator of cardiovascular and cerebrovascular disease. Monocyte- and macrophage-related inflammation rather than T cell activation is regarded as a major cause of chronic inflammation in the context of treated HIV infection^4–7^.

Circulating monocytes can differentiate into macrophages and have distinct morphologies, transcriptomes, and biological functions. The bulk RNA sequencing and microarray revealed that monocytes isolated from healthy donors’ blood express high levels of chemokines and the corresponding receptors to regulate inflammation. In contrast, the prominent genes in macrophages differentiated from monocytes are related to metabolic processes and the phagocytic capacity, which is involved in the eradication of various pathogens and cell debris and metabolizing lipid, steroid, fatty acid, and amino acid^8–10^. Therefore, while there are similarities between monocytes and macrophages, there are also important differences that hold true for the Dll4-Notch pathway investigated in this report.

Notch family comprises highly conserved trans-membrane proteins, including receptors (Notch1-Notch4) and ligands (Delta-like ligand 1 (Dll1), Delta-like ligand 4 (Dll4) and Jagged 1 (Jag1). The ligands bind to Notch receptors of adjacent cells, triggering proteolytic cleavage of Notch by γ-secretase, to release the Notch intracellular domain (NICD). NICD translocates into the nucleus and promotes downstream gene transcription ^11, 12^. Dll4 is usually restricted in endothelial cells (ECs). Dll4^-/-^ and Dll4^-/+^ mice are embryonic lethal and show severe and precocious vascular defects, demonstrating the pivotal role of Dll4 in vascular development^13^.

Recently, increased evidence showed the role of Dll4 in macrophage polarization and atherosclerosis. Cytokines including lipopolysaccharides (LPS) and interleukin (IL-1β) can induce Dll4 expression in macrophages in vitro through nuclear factor-κB (NF-κB) and Notch-dependent pathways^14, 15^. Enhanced Dll4 subsequently activates Notch signaling to lead to proinflammatory effects indicated by enhanced mRNA levels of TNFα, MCP1, iNOS, IL-6, and IL-12^16–18^. Dll4 inhibits the upregulation of IL-4 induced M2 markers such as CD206 and CD200R^19^. Survival of macrophages upon M2 polarization was also strongly reduced in the presence of Dll4. Dll4 induces caspase3/7-dependent apoptosis during M2 but not M1 macrophage polarization^15, 20–22^, which contributes to the pathogenesis of atherosclerosis^18^. In PLWH, fecal microbial translocation from the gastrointestinal (GI) tract to the systemic circulation, causes a significant increase in plasma LPS^23–26^ and LPS is the strongest stimulus for Dll4 expression in macrophages^15^. However, the role of Dll4 in the monocytes of PLWH has not been explored. We hypothesize that expression of Dll4 and subsequent Dll4-Notch signaling activation in monocytes result in the release of cytokines/chemokines from circulating monocytes, contributing to the persistent chronic systemic inflammation in PLWH and promoting HIV-associated comorbidities. This cytokine-Dll4-Notch-cytokine positive feedback loop could be a novel therapeutic target for HIV related CVD.

## Methods

All supporting data are available within the article.

### Human subjects

Study procedures are fully described in previous publication^27^. As a part of an ongoing study approved by the Institutional Research Subjects Review Board (RSRB) at the University of Rochester, a total of 105 HIV-infected (HIV) men and women on cART treatment (mean ± SD age = 53.2 ± 11.2 years) and 105 HIV-uninfected (non-HIV) age and sex matched controls (mean ± SD age = 51.0 ± 16.4 years) were evaluated in this study. Given the sex distribution of HIV infection in our clinics, more men than women were enrolled. Blood samples were acquired after written informed consent. Total 210 patients’ plasma and PBMC samples were aliquoted and cryopreserved at liquid nitrogen for no more than two years for plasma exDll4 measurement and monocyte subsets analysis, respectively.

For co-enrollment to pursue additional experiments, 15 non-HIV (N=9, male; N=6, female) and HIV patients (N=22, male; N=6, female) were recruited. In this subset of participants, approximately 20 ml of whole blood was collected in K2 EDTA vacutainers (4 vacutainers) and processed within 2h of collection. Plasma and PBMCs from both cohorts were studied. According to the manufacturer’s manuals, plasma, PBMC, and granulocytes were isolated from peripheral blood using Ficoll-Paque® Plus (GE Healthcare). Briefly, whole blood was gently transferred on the Ficoll-Paque® Plus, after spinning down at 400g for 40 minutes at room temperature, plasma from the upper layer, PBMC from the intermediate layer, and granulocytes from the lower layer were collected. PBMCs and granulocytes were cryopreserved in CryoStor® CS10 (#100-1061, StemcCell Technologies), which can provide maximum post-thaw cell recovery and viability.

### Cell culture

THP1, a monocyte cell line isolated from peripheral blood from a male acute monocytic leukemia patient, was used for the experiments (#TIB-202, ATCC). To induce Dll4 expression, THP1 cells culture media was treated with vehicle, 100ng/mL of LPS (#L3012, Sigma-Aldrich), or 10ng/mL of recombinant human TNF-α (#300-01A, PeproTech) for 6, 12, 24, and 48h. mRNA and protein were harvested for qPCR and western blot, respectively. Culture supernatant (CS) was harvested for Dll4 ELISA assay (#OKEH02321, Aviva Systems Biology) to detect the extracellular domain of Dll4, according to the manufacturer’s instructions. To generate macrophage and detect Dll4 expression, THP1 cells were stimulated with phorbol 12-myristate 13-acetate (PMA, 100ng/ml) for 24h, followed by LPS or TNFα for another 48h. THP1 cells were transfected with control or Dll4 siRNA (Sigma-Aldrich) at 10 µM by Lipofectamine™ RNAiMAX (#13778, Invtrogen), or culture media was treated with vehicle or DAPT (N-[N-(3, 5-difluorophenacetyl)-l-alanyl]-s-phenylglycinet-butyl ester, D5942, Sigma-Aldrich) at 10µM for 24h in the presence of LPS or TNFα stimulation. To inhibit NF-κB signaling, THP1 cells were treated with control or JSH-23 (#J4455, Sigma-Aldrich) at 10 µM for 48h in the presence of TNFα stimulation.

### Flow cytometry

Frozen PBMCs were quickly thawed in a 37°C water bath and diluted ten times with a warmed completed monocyte culture medium. After spinning down, the supernatant was removed carefully without affecting the cell pellet. To reduce aggregation of concentrated and cryopreserved cells suspension following thawing, we resuspended cells in 0.1 mg/mL of DNase I solution (#07900, StemcCell Technologies) and incubated them at room temperature for 15 minutes. We filtered aggregated suspension through a 40µm cell strainer before staining for optimal cell separation. After lysing red blood cells and blocking Fc-receptors, to detect the expression of membrane bound Dll4 by flow cytometry, human PBMCs were stained with anti-CD14-APC (1:100), anti-CD16-FITC (1:100), and anti-Dll4-PE (1:20) (BioLegend) and incubated on ice for 30 minutes in the dark. In the end, cells were washed three times to remove any unbound antibodies and resuspended in 2% formaldehyde/PBS and analyzed on a flow cytometer. FlowJo™ Software was used for single-cell flow cytometry analysis.

### ELISA

Human extracellular Dll4 (aa27-302) was measured using a Quantikine ELISA kit from Aviva Systems Biology (OKEH02321) following the manufacturer’s instructions. Dll4 antigen was detected following the protocol. The blank-corrected absorbances of the standard curve measurements indicated that this kit has a high sensitivity to Dll4 detection. We applied 100ul of patient plasma without dilution for ELISA analysis, and duplicate wells for each sample.

### qRT-PCR and immunoblotting

We performed qRT-PCR and western blots as previously described^28, 29^. Results were normalized by setting the densitometry of control samples to 1.0. Commercial primary antibodies used for western blot were anti-human Dll4 (#2589, CST), cleaved Notch1 (#4147, CST), Notch1 (#4380, CST), Notch2 (#5732, CST), Notch3 (ab23426, Abcam), Dll1 (sc-377310, Santa Cruz Biotechnology), Jag1 (#2620, CST), β-actin (PA1-183, Thermo Fisher Scientific) and GAPDH (MAB374, Millipore).

### Statistical analysis

Statistical analyses were performed using GraphPad Prism Statistical Analysis software and SAS 9.4 (SAS Institute Inc, Cary, NC). All values are expressed as mean ± SEM from three to eight samples. Bivariate analyses of association between HIV status and demographics and clinical variables were performed using the Student’s *t*-test or its nonparametric analogy of Wilcoxon Rank-sum test for continuous variables, and Chi-square test or its nonparametric version of Fisher exact test for categorical variables. Pearson and Spearman correlation analysis were used to test the association between two continuous variables. Multiple regression was used to explore the heterogeneous association between exDll4 and monocyte subpopulation distributions depending on gender. For all analyses, a *P* value < 0.05 was considered significant.

## Results

### Proinflammatory stimuli increase Dll4 expression and secretion and Notch1 activation in monocytes

LPS and TNFα are significantly increased in the plasma of HIV patients^23–26^. Aikawa’s group showed cytokines increase Dll4 expression in human macrophages, which is critical for atherosclerosis^15^. However, monocytes have different transcriptomes and biological functions compared with macrophages and the effect of inflammatory factors on Dll4 expression in monocytes has not been previously investigated. Given that monocytes can differentiate into macrophages, we hypothesized that LPS and TNFα would induce Dll4 expression in monocytes. Because the average life span of monocytes is 2-6 days in human blood and 2 days in mouse blood^30^, THP1 cells were treated with LPS (100ng/mL) or TNFα (10ng/mL) for 6, 12, 24, and 48h. Without proinflammatory stimuli, expression of Dll4 was extremely low in THP1 cells. After stimulation, we observed a significant time-dependent increase in Dll4 mRNA and protein expression. The peak of Dll4 mRNA was at 24h (increased 35 and 17.1-fold by LPS and TNFα respectively, Figure 1A), and the peak Dll4 protein expression was at 48h (increased 71.1 and 23.6 folds by LPS and TNFα respectively, Figure 1B). Ligand Dll1 expression was comparable and Jag1 expression was hard to detect before and after cytokine treatments (Figure S1A-S1B). Elevated Dll4 could activate canonical Notch signaling. Thus, the cleavage of Notch receptors was detected in monocytes. Consistent with the enhanced Dll4 expression, Notch1 activation (detected by specific N1-ICD antibody, Val1744) increased 124.5 and 21.5 folds in response to LPS and TNFα, respectively. While Notch2 activation was not (measured by Notch2-NTM antibody, Figure S1A-S1B) compared to the control group. Notch 3 and 4 expression was not detectable (data not shown).

**Figure 1.**
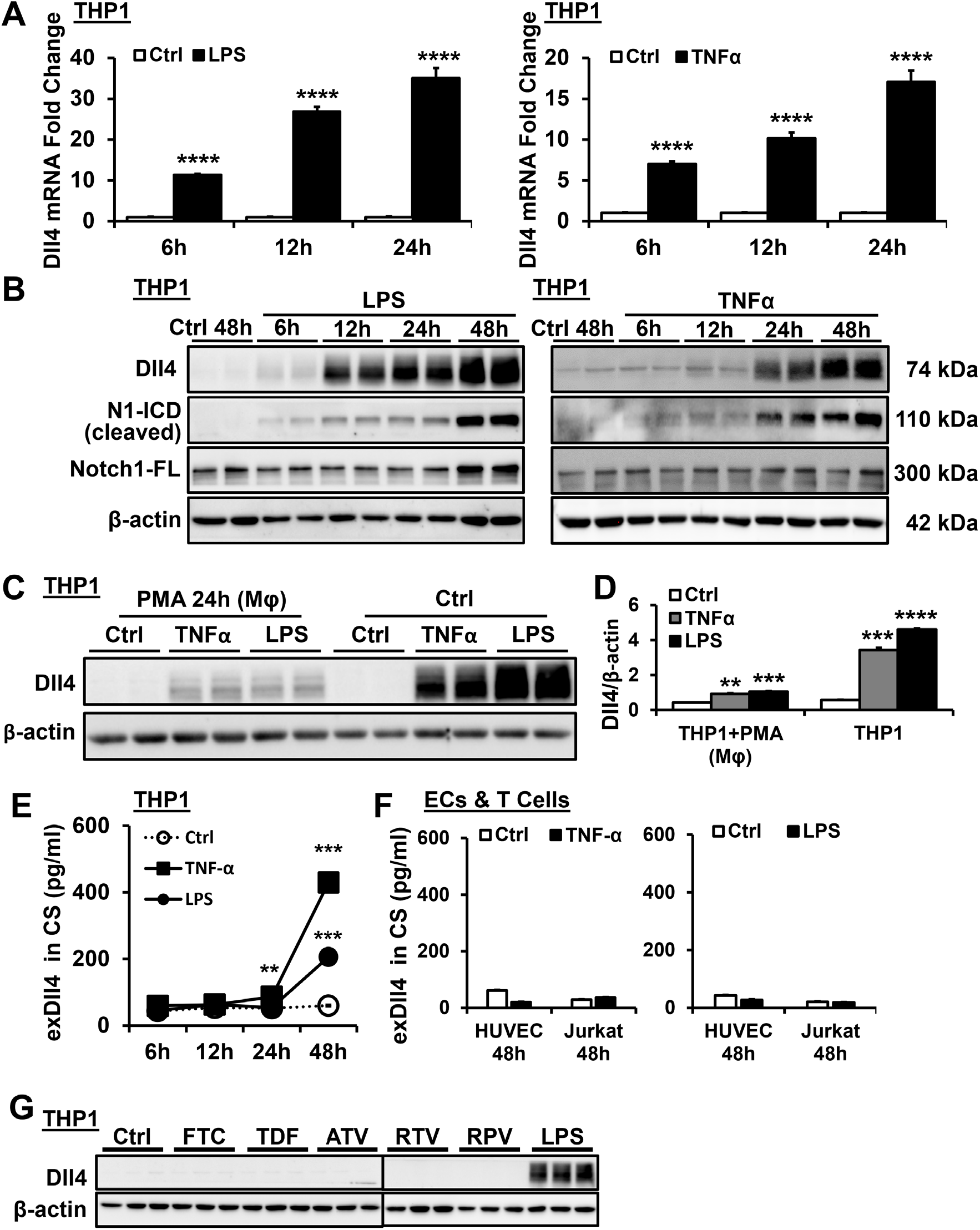
LPS and TNFα increase Dll4 expression and secretion and Notch1 activation in monocytes. **(A)** qPCR of Dll4 and **(B)** western blot of Dll4, N1-ICD and Notch1-full length (FL) in THP1 cells with or without LPS or TNFα stimulation at 6, 12, 24 and 48h (n=6). **(C-D)** Western blot of Dll4 in THP1 cells and macrophages with or without LPS or TNFα stimulation at 48h (n=4) and quantification. **(E-F)** Dll4 levels in CS of THP1, HUVEC and Jurkat after LPS or TNFα stimulation were detected by ELISA (n=6-12). **(G)** Western blot of Dll4 in THP1 cells treated with or without anti-HIV drugs at 48h (n=4). Data shown as Mean±SEM; *P* values were calculated using the Student’s *t*-test, and \*\**P*<0.01; ***, *P*˂0.001; ****, *P*˂0.0001 compared with control group. CS, culture supernatant. exDll4, extracellular Dll4. Mφ, macrophage.

To compare the Dll4 levels in response to cytokines in monocytes and macrophages, THP1 cells were differentiated into macrophages by stimulating with phorbol 12-myristate 13-acetate (PMA) or vehicle for 24h, then stimulated with cytokines. LPS or TNFα administration for 48h significantly induced Dll4 protein expression in both macrophages and monocytes, however, the increase in monocytes was 4.4 folds and 3.7 folds higher than that in macrophages respectively (Figure 1C-1D), suggesting that excessive Dll4 in monocytes may play an important role on systemic inflammation.

Monocytes can release cell surface proteins (especially proteins in small size) by phagocytosis^31^. Therefore, we postulate that monocytes can secrete Dll4. THP1 cells were stimulated with LPS or TNFα at various time points, and culture supernatant (CS) was harvested for ELISA assay to detect extracellular Dll4 (exDll4). We found that exDll4 levels in CS increased by 188.4% by LPS at 48h, 53.8%, and 440.5% by TNFα at 24 and 48h respectively compared with the control groups (Figure 1E). However, CS exDll4 levels were consistently low in human umbilical cord endothelial cells (HUVECs) and Jurkat cells (T cell cell line) with/without LPS or TNFα administration (Figure 1F), suggesting that monocytes are the major source of exDll4. Anti-HIV drugs increase inflammation in circulating monocytes^4^, thus we measured the Dll4 expression in THP1 cells after administration of anti-HIV drugs. We found that these treatments had no effect on Dll4 expression, excluding the role of cART in Dll4 expression in monocytes and secretion from monocytes (Figure 1G). The data implies that inflammatory factors are critical for Dll4 expression and Notch1 activation in monocytes and exDll4 release from monocytes.

### Dll4-Notch1 signaling activation enhances inflammation in monocytes

Dll4-Notch signaling leads to inflammation in macrophages^16–18^, we speculated similar phenomena will be observed in monocytes. Therefore, the gene expression of both Notch target genes and proinflammatory cytokines/chemokines was measured by qPCR. As expected, LPS or TNFα treatment for 24h induced a robust increase of Notch target genes, including Hes1, Hes2, Hes7, Hey1, and Hey2 but did not affect Hes4 and Hes6, as well as increased cytokines/chemokines, such as IL-6, IL-1β and TNFα mRNA (Figure 2B-2C, 2E-2F). To determine the role of Dll4-Notch1 signaling in monocyte inflammation, we knocked down Dll4 by siRNA or inhibited Notch activation by a γ-secretase inhibitor DAPT (N-[N-(3, 5-difluorophenacetyl)-l-alanyl]-s-phenylglycinet-butyl ester) in THP1 cells for 24h in the presence of LPS or TNFα. Silencing of Dll4 or inhibition of Nocth1 activation remarkably suppressed LPS or TNFα -induced Notch target gene and proinflammatory gene expression (Figure 2B-2C, 2E-2F). Dll4 siRNA reduced Dll4 and N1-ICD protein expression in monocyte (Figure 2A and 2D). Interestingly, although DAPT inhibited Notch1 activation, it did not affect Dll4 mRNA and protein expression (Figure 2D-2E) which is opposite to our and other groups’ findings of Dll4-Notch-Dll4 positive feedback loop in endothelial cells (ECs) and macrophages^15, 32^. The different effects of Dll4 siRNA and DAPT on Dll4 expression are possible due to the different transcriptomes of ECs, monocytes, and macrophages^10^. All the findings above indicated that Dll4-Notch1 signaling enhances inflammation in monocytes.

**Figure 2.**
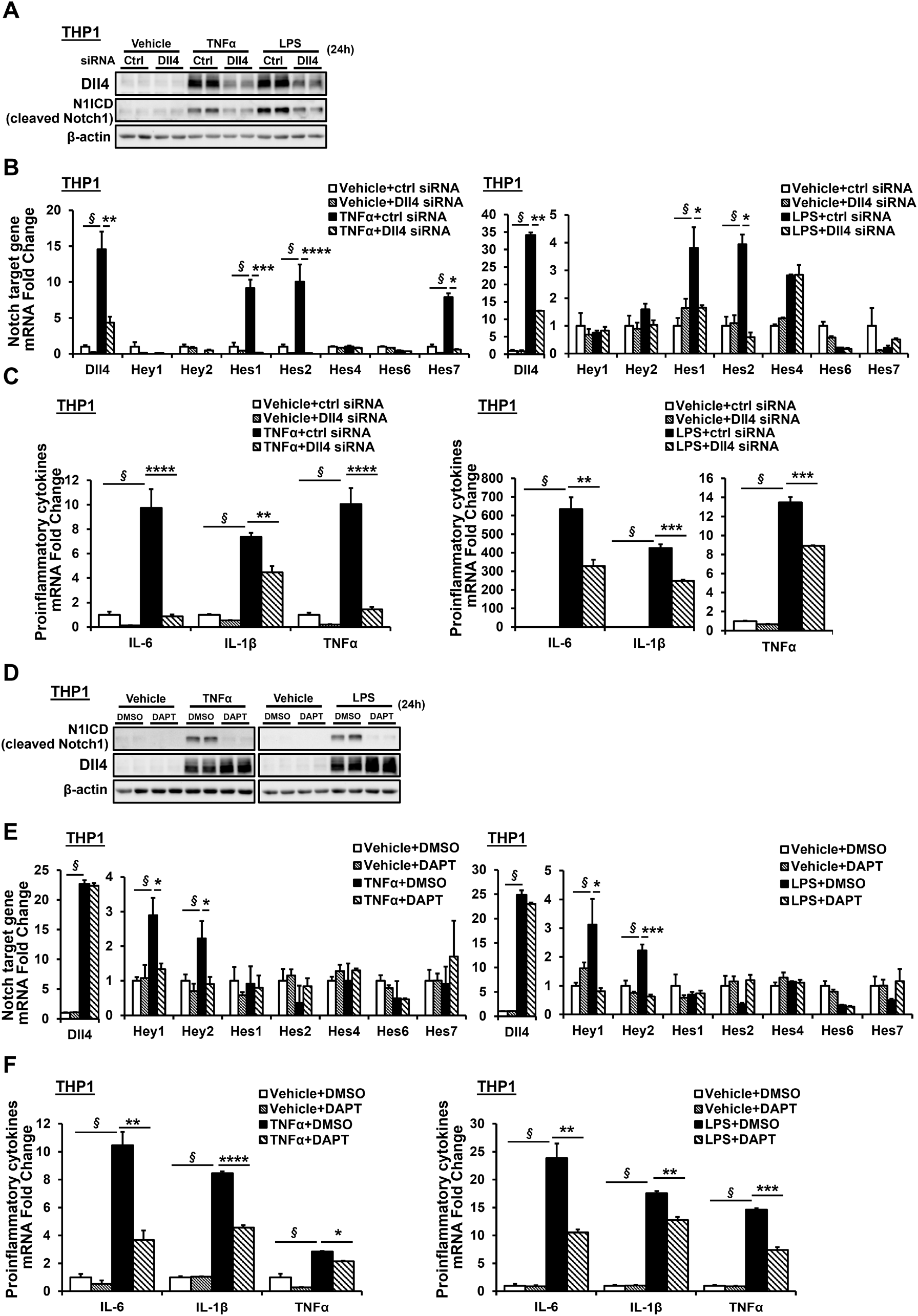
Dll4-Notch1 signaling activation enhances inflammation in monocytes. **(A-C)** After Dll4 silencing or **(D-F)** inhibition of Notch signaling, **(A, D)** western blot of Dll4, N1-ICD and Notch1-full length (FL) and **(B-C, E-F)** qPCR of Notch target genes and pro-inflammatory genes mRNA expression in THP1 cells with or without LPS or TNFα stimulation at 24h (n=8). Data shown as Mean±SEM; *P* values were calculated using the Student’s *t*-test, and \**P*<0.05; \*\**P*<0.01; ***, *P*˂0.001; ****, *P*˂0.0001 compared with control group. § indicates significant difference between vehicle and cytokine treated control siRNA or DMSO groups.

### Enhanced membrane-bound Dll4 (mDll4) in monocytes and elevated plasma exDll4 in PLWH

Chronic inflammation is critical for cardiovascular diseases in PLWH thus we speculated an elevation of exDll4 in plasma and increased mDll4 and Notch target genes in monocytes of PLWH. We obtained PBMCs and plasma samples of PLWH under cART treatment and age matched non-HIV patients. Clinical and demographic characteristics are presented in Table 1. PBMCs were isolated from the blood of the aforementioned cohort^27^ and incubated with CD14 and CD16 (monocyte specific markers), and Dll4 antibodies for flow cytometry analysis. Compared with non-HIV patients, PLWH had a 1.9-fold significant elevation of the intensity of mDll4 in monocytes and a 2.1-fold increase in the percentage of mDll4 positive monocytes (Figure 3A and 3E). Considering the potential sex effects, we analyzed the data from males and females separately, and observed similar results to combined data (Figure 3B and 3F). Human monocytes are classified into three subsets: the classical, intermediate, and nonclassical monocytes^33^. Classical monocytes sequentially give rise to intermediate and then to non-classical monocytes^30^. Classical monocytes comprise most of the circulating monocytes and confer anti-inflammatory responses^34, 35^. In contrast, intermediate and non-classical monocytes have been considered pro-inflammatory monocyte subsets in infectious and inflammatory conditions (e.g., sepsis and HIV)^33, 34, 36, 37^. Therefore, we also analyzed mDll4 expression in three subsets of monocytes. Compared to the sex matched non-HIV controls, the intensity of mDll4 was significantly elevated by 177.7% in the classical subset, 113.8% in the intermediate subset and 96.8% in the non-classical subset of HIV male patients, and increased by 89.2% in classical, 81.5% in intermediate subsets of HIV female patients (Figure 3C-3D). While no differences were observed in the non-classical subset between HIV and non-HIV female patients (Figure 3D). Notably, the intermediate subset had the highest mDll4 intensity in both genders of HIV patients. In contrast, the percentage of mDll4 positive classical monocytes significantly increased by 150.5% and 179.65% in HIV male and female patients respectively. The percentage of mDll4 positive intermediate monocytes increased by 732.7% in HIV female patients (Figure 3G-3H). This analysis suggested that mDll4 was expressed at higher levels in all three subsets of monocytes in male PLWH and in the classical and intermediate subsets of monocytes in female PLWH, compared to HIV uninfected individuals. Of notice female PLWH intermediate monocytes had the highest mDll4 intensities. The different patterns of Dll4 intensity vs percentage in intermediate subpopulations imply specific regulatory mechanisms of Dll4 expression. Moreover, mDll4 was hardly measured (data not shown) in non-monocyte PBMCs (including T cells, B cells, and NK cells) and was not detectable in the granulocytes of HIV patients (Figure 3I).

**Figure 3.**
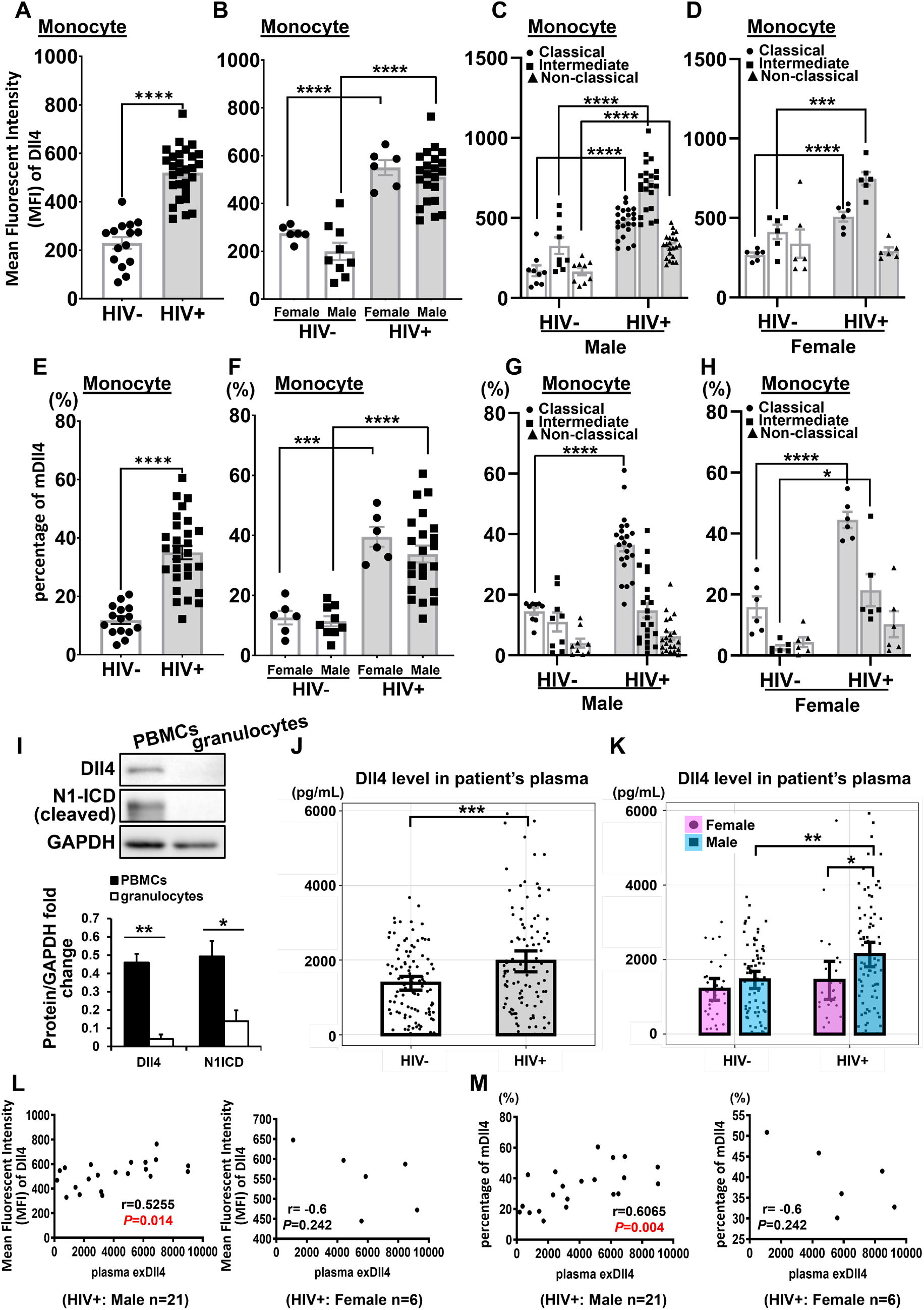
Elevated mDll4 in monocytes and enhanced plasma exDll4 in PLWH. **(A-D)** Intensity and **(E-H)** percentage of mDll4 in monocytes of HIV patients (n=22, male; n=6, female) compared with non-HIV patients (n=9, male; n=6, female). **(I)** Western blot of Dll4 and N1-ICD protein expression in PBMCs and granulocytes of HIV patient (n=3). **(J-K)** exDll4 levels in plasma of HIV and non-HIV patients (n=105 per group). Spearman correlation analysis between plasma exDll4 and **(L)** intensity and **(M)** percentage of mDll4 in monocytes in HIV patients (n=21, male; n=6, female). Data shown as Mean±SEM; *P* values were calculated using the Student’s *t*-test, and \**P*<0.05; \*\**P*<0.01; ***, *P*˂0.001; ****, *P*˂0.0001 compared with the control group.

**Table 1.**
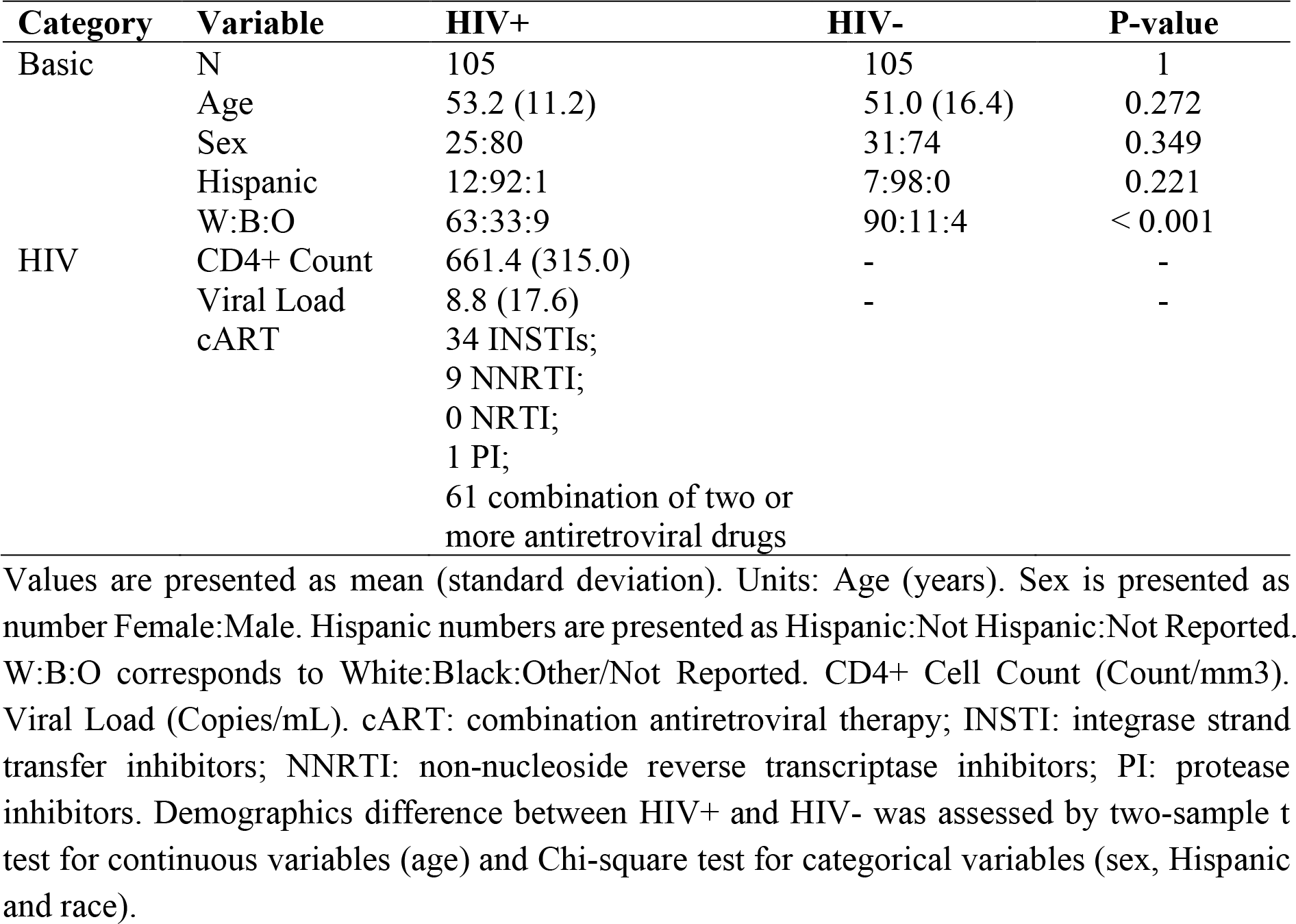
Clinical and demographic characteristics of the patients.

| Category | Variable | HIV+ | HIV- | P-value |
| --- | --- | --- | --- | --- |
| Basic | N | 105 | 105 | 1 |
|  | Age | 53.2 (11.2) | 51.0 (16.4) | 0.272 |
|  | Sex | 25:80 | 31:74 | 0.349 |
|  | Hispanic | 12:92:1 | 7:98:0 | 0.221 |
|  | W:B:O | 63:33:9 | 90:11:4 | < 0.001 |
| HIV | CD4+ Count | 661.4 (315.0) | - | - |
|  | Viral Load | 8.8 (17.6) | - | - |
|  | cART | 34 INSTIs; |  |  |
|  |  | 9 NNRTI; |  |  |
|  |  | 0 NRTI; |  |  |
|  |  | 1 PI; |  |  |
|  |  | 61 combination of two or more antiretroviral drugs |  |  |

Values are presented as mean (standard deviation). Units: Age (years). Sex is presented as number Female:Male. Hispanic numbers are presented as Hispanic:Not Hispanic:Not Reported. W:B:O corresponds to White:Black:Other/Not Reported. CD4+ Cell Count (Count/mm<sup>3</sup>). Viral Load (Copies/mL). cART: combination antiretroviral therapy; INSTI: integrase strand transfer inhibitors; NNRTI: non-nucleoside reverse transcriptase inhibitors; PI: protease inhibitors. Demographics difference between HIV+ and HIV- was assessed by two-sample t test for continuous variables (age) and Chi-square test for categorical variables (sex, Hispanic and race).

To determine the released exDll4 in circulation, ELISA was performed using plasma samples of cART treated PLWH (N=80, male; N=25, female, mean age 53) and age matched non-HIV patients (N=74, male; N=31, female, mean age 51). Compared with non-HIV patients, plasma exDll4 levels were significantly elevated by 44.5% in PLWH (Figure 3J). To explore the potential impact of sex differences, we also separated the data from males and females. No significant difference was observed between non-HIV and HIV female subjects. However, plasma levels of exDll4 were increased by 54.84% in male PLWH compared with the non-HIV male controls (Figure 3K). Circulating exDll4 was comparable between non-HIV males and females, while exDll4 of male PLWH was increased by 59.6% compared to female PLWH. Moreover, the Spearman correlation test showed that plasma exDll4 was positively associated with Dll4 intensity of monocyte (Figure 3L, r=0.5255, *P*<0.05) or the percentage of Dll4 positive monocytes (Figure 3M, r=0.6065, *P*<0.01) in male PLWH, suggesting that exDll4 was derived from circulating monocytes. Although mDll4 was highly expressed in monocytes of PLWH irrespective of sex, increased plasma levels of exDll4 were only observed in male PLWH compared to uninfected controls. This implies that different mechanisms of the monocyte secretome are at play in males and females in response to inflammatory stimuli.

### Elevated monocyte Dll4-Notch1 signaling and inflammatory gene expression in PBMCs of PLWH

To further determine the Dll4-Notch1 signaling activation in PLWH PBMCs, we measured Notch targeted gene expression (hey1/2, hes1-7) in isolated PBMCs using qPCR. Because mDll4 was comparable in monocytes of non-HIV male and female patients, we used the result of all non-HIV patients as the control. Consistent with enhanced mDll4 in monocytes, Hey1, Hes1, and Hes7 mRNA expression in male PLWH increased 2.48, 2.16, and 3.28-fold respectively compared with the non-HIV controls, while Hey2, Hes2, Hes4, and Hes6 did not change and Hes5 was not detectable (Figure 4A). In contrast, only Hes1 expression enhanced in females PLWH, and Hes7 expression decreased, suggesting the different Notch target genes between males and females under inflammatory conditions. We also measured the proinflammatory cytokine/chemokine gene expression in PBMCs by qPCR. IL-6, IL-1β, and TNFα expression increased 14.2, 30.6, and 2.5-fold respectively in PBMCs of male PLWH and elevated 4.1, 14.1, and 2.6-fold in female PLWH (Figure 4B). Our findings indicate increased activation of Dll4-Notch1 signaling and inflammation in monocytes of HIV patients.

**Figure 4.**
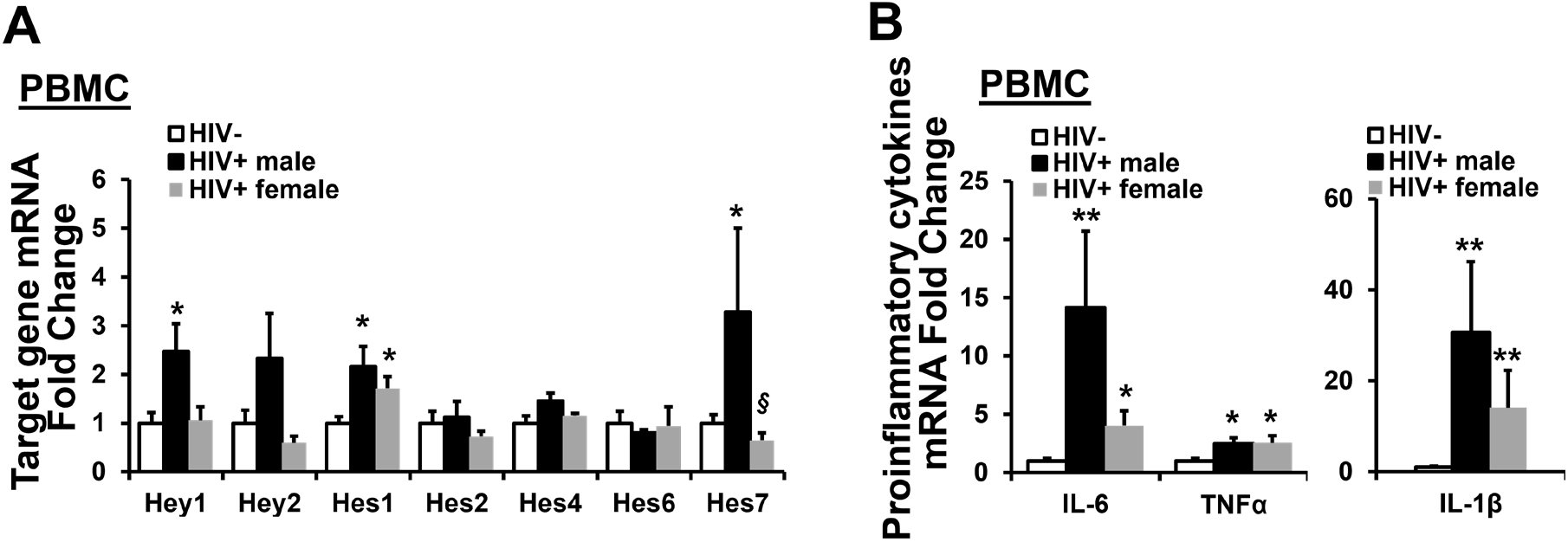
Increased Dll4-Notch1 signaling and inflammatory gene expression in monocytes of PLWH. **(A)** Notch target genes and **(B)** pro-inflammatory genes mRNA expression in PBMCs of HIV (n=5, male; n=6, female) and non-HIV patients (n=2, male; n=4, female). Data shown as Mean±SEM; *P* values were calculated using the Student’s *t*-test, and \**P*<0.05; \*\**P*<0.01 compared with control group. § indicates significant difference between male and female HIV patients.

### Plasma exDll4 is associated with inflammation in male PLWH

Given that Dll4 promotes M1 but inhibits M2 macrophage polarization^22^, we hypothesized that elevated exDll4 might be associated with monocyte subpopulation distribution. We observed that plasma exDll4 was negatively associated with classical monocytes (Figure 5A, *β* =-0.0023, *P*<0.05) and positively associated with proinflammatory monocytes including intermediate and non-classical monocytes (Figure 5B, *β* =0.0021, *P*<0.05). The association was only observed in male PLWH but not in female patients.

**Figure 5.**
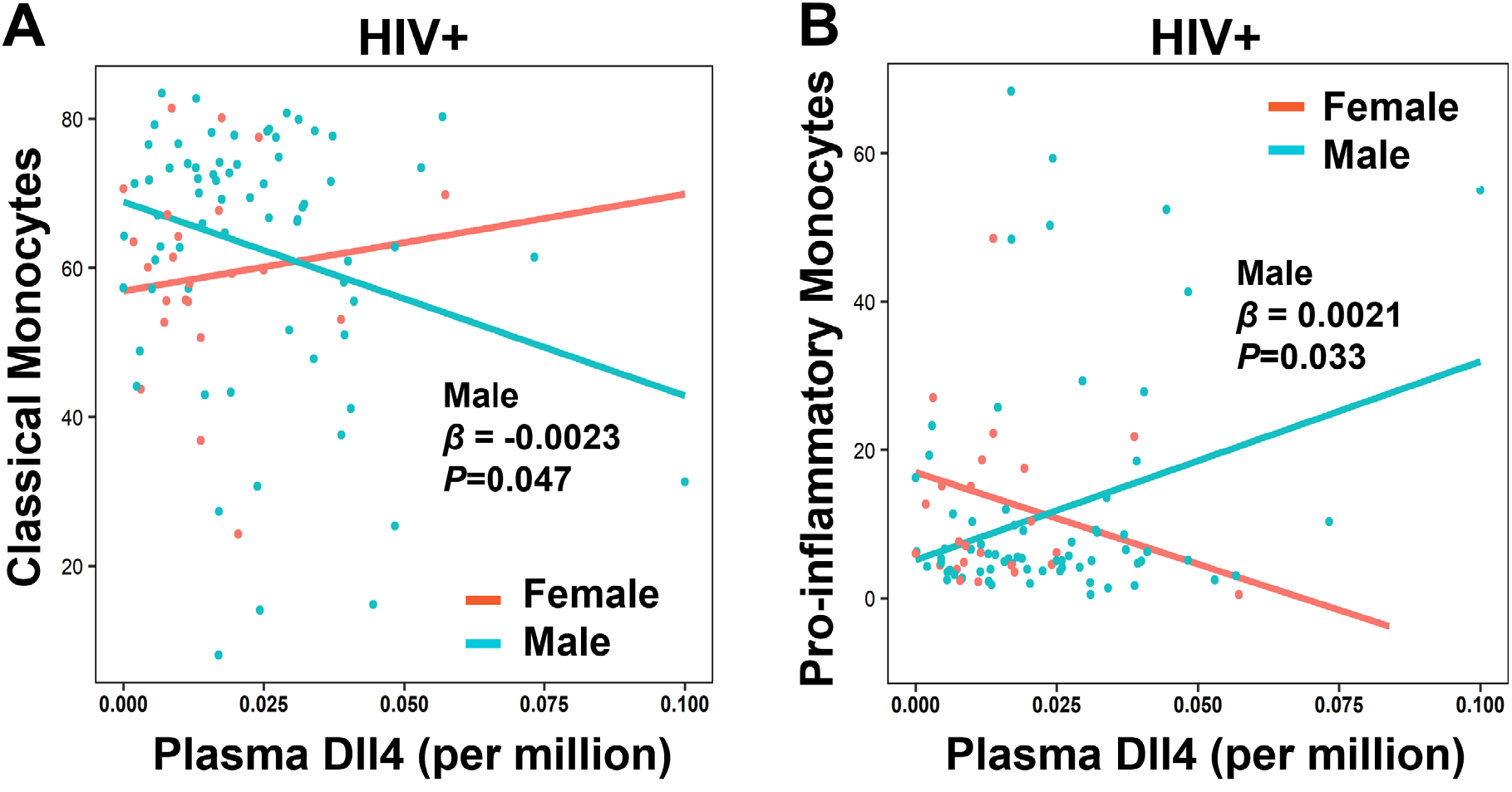
Plasma exDll4 is associated with inflammation in male PLWH. **(A)** Classical monocytes and **(B)** pro-inflammatory monocytes (intermediate and non-classical monocytes) associated with plasma exDll4 in PLWH by Pearson correlation analysis. Age and HIV status were adjusted. Female=50; male =124. *β*, slopes.

## Discussion

The major finding of this study is that monocyte Dll4 expression is a potential contributor to persistent systemic inflammation in PLWH. In vitro experiments show that pro-inflammatory factors can trigger Dll4 expression and Dll4-Notch1 signaling activation in monocytes, induce inflammatory gene expression, and exDll4 secretion. Dll4 silencing and inhibition of Notch activation diminish the LPS or TNFα -induced inflammation. Inflammatory factors cause exDll4 release from monocytes but no other cell types. In vivo, we observe increased expression of mDll4 and inflammatory genes in monocytes derived from PLWH compared to uninfected individuals. Surprisingly, elevated plasma exDll4 is only observed in male PLWH, which also showed a positive correlation with proinflammatory monocytes.

This sex-based difference in plasma levels of exDII4 requires further investigations. The difference between PLWH males and females is possibly due to the differences in the monocyte transcriptome and monocytic cell secretome of male and female patients, perhaps caused by different patterns of Notch target genes serving as transcription factors. For example, Hes1 was enhanced in monocytes of both male and female patients, while Hes7 increased about 3 folds in HIV male patients but significantly decreased in HIV female patients compared with non-HIV patients. This concept is supported by several studies on patients with chronic low-grade inflammation, which demonstrated that males had a stronger cytokine production response to LPS^38–40^. Our findings are consistent with the previous literature suggesting that the immune response to pathogens or self-antigen differs in males and females^41^. This observation applies also to circulating monocytes in the context cardiovascular risk factors such as obesity^42^. Our findings may provide potential mechanisms for why men have a higher prevalence of CVD than women in HIV patients^43^. Our findings imply that while mDll4 is a potential biomarker of systemic inflammation for both genders, exDll4 is more useful in males. Given the pivotal role of systemic inflammation in HIV related cardiovascular diseases, mDll4 is a potentially promising therapeutic target.

Ectopic expression of Dll4 in monocytes contributes to systemic inflammation in HIV patients. In vitro evidence shows that inflammatory molecules, such as LPS, IL-1β or minimally-modified low-density lipoprotein but not TNFα, increase Dll4 expression in macrophages through nuclear factor-κB (NF-κB) and Notch signaling^15^. Increased expression of Dll4 activates Notch to trigger inflammatory responses in macrophages ^15–18^. In contrast, we demonstrate that LPS and TNFα but not IL-1β (data not shown), the major regulators of inflammatory responses, can promote Dll4 expression in monocytes via the NF-κB pathway (Figure S2A-S2B). Elevated Dll4 expression was independent of Notch signaling (Figure 2D), which contradicts the Dll4-Notch-Dll4 positive feedback loop in endothelial cells (ECs)^32^ and macrophages^15^, implying a distinct regulatory mechanism of Dll4 expression in monocytes. In addition, cytokines induced Dll4 protein expression was 3-4 folds higher in monocytes than in macrophages. Dll4-activated Notch1 signaling augmented typical peripheral levels of proinflammatory cytokines IL-1β, IL-6, and TNFα in monocytes, while Dll4 and Jag1 mediated Notch signaling (Jag1 is highly expressed in macrophages but extremely low in monocytes, Figure S1) induced iNOS, PTX3, and Id1etc in macrophages to promote atherosclerosis progression^15^. Thus, our results suggest that increased Dll4-Notch signaling in monocytes promotes the expression of inflammatory factors and the cytokine-Dll4-Notch-cytokine positive feedback loop propagates these effects leading to persistent inflammation in HIV patients, which exacerbates CVD.

Resembling our findings in vitro, our clinical results revealed a significant increase of mDll4 expression in monocytes, Notch target genes, and pro-inflammatory genes expression in PBMCs of HIV patients. We further proved that mDll4 was only induced in monocytes but not T cells, B cells, or neutrophils in PLWH. All these findings further support the substantial contribution of monocyte Dll4 contribution in sustained inflammation in PLWH.

Our study demonstrates that exDll4 is a potential inflammatory marker in male PLWH. In vitro, we found that proinflammatory stimuli induced secretion of exDll4 in the culture supernatant of monocyte (THP1 cell lines generated from a male acute monocytic leukemia patient) but not in ECs and T cells. In vivo, elevated plasma exDll4 was observed in male PLWH. Consistent with our finding, in patients infected with mycobacterium tuberculosis, circulating Dll4 was significantly increased^44, 45^. However, these previous studies did not answer why exDll4 was increased in patients infected with tuberculosis nor did they address the origin of the increased exDll4 in the serum samples. Our study is the first to demonstrate that increased exDll4 was released from monocyte exposed to proinflammatory stimuli in PLWH.

Our study has several limitations. First, the relatively small sample size of female PLWH. These results will need to be repeated in a larger cohort of female patients to confirm the findings. Second, due to the limitation of monocyte cell numbers from patients, we could only detect gene expression in PBMCs, not monocytes. Third, the functions of exDll4 need to be further investigated.

In summary, we have demonstrated that pro-inflammatory stimuli trigger Dll4-Notch1 signaling activation in monocytes, exDll4 secretion, and enhances monocyte pro-inflammatory phenotype, contributing to persistent systemic inflammation in HIV patients. mDll4 in monocyte and plasma exDll4 could be potential biomarkers of HIV-related CVD caused by monocyte-mediated inflammation. Our findings implicate that specifically targeting Dll4 in monocyte could be a novel therapeutic strategy for CVD.

## Sources of Funding

This work was supported by Jinjiang Pang’s grants from the National Institutes of Health (R01 HL122777-08, R01 HL122777-08S1), American Heart Association Innovative Project Award (19IPLOI34760446), and the Vaccine and Treatment Evaluation Unit (VTEU) pilot grant of University of Rochester. The HIV cohort was supported by R01 AG054328 (Sanjay Maggirwar and Giovanni Schifitto (MPIs) and R01MH118020 (Giovanni Schifitto, PI).

## Conflict of Interest Statement

None.

## Novelty and Significance

### What Is Known?

- Persistent systemic inflammation is a driving force for the progression of cardiovascular and cerebrovascular diseases in people living with HIV.
- Monocyte- and macrophage-related inflammation rather than T cell activation is a major cause of chronic inflammation.

### What New Information Does This Article Contribute?

- Pro-inflammatory stimuli trigger Dll4 expression and Dll4-Notch1 signaling activation in monocytes, induce inflammatory gene expression, and exDll4 secretion.
- Enhanced mDll4 expression and inflammatory genes in monocytes are consistently observed in both HIV male and female patients.
- Elevated plasma exDll4, which shows a positive association with proinflammatory monocytes, was only observed in male PLWH, implying sex differences in Dll4 secretion.
- mDll4 in monocyte and plasma exDll4 could be potential biomarkers of HIV-associated CVD. Targeting monocyte Dll4 could be a novel therapeutic strategy for HIV-associated CVD.

## Summary

In HIV patients under combination antiretroviral therapy (cART), persistent systemic inflammation is a driving force for the progression of comorbidities, such as cardiovascular diseases (CVD). Monocyte and macrophage-related inflammation rather than T cell activation is a major cause of chronic inflammation. However, the underlying mechanism of how monocytes cause persistent systemic inflammation in HIV patients is elusive. We demonstrated that pro-inflammatory stimuli LPS and TNFα trigger Dll4 expression and Dll4-Notch1 signaling activation in monocytes, induce inflammatory gene expression, and Dll4 secretion (extracellular Dll4, exDll4). Monocytes are the major source of exDll4. Cytokines-induced Dll4 protein expression was 3-4 folds higher in monocytes than in macrophages, indicating an essential role of monocyte Dll4 in systemic inflammation compared to the role of Dll4 in macrophages in atherosclerotic plaques. Enhanced Dll4 expression and inflammatory genes in monocytes are consistently observed in both HIV male and female patients. Elevated plasma exDll4, which shows a strong positive association with proinflammatory monocytes, was only observed in HIV male patients, implying sex differences in Dll4 secretion in HIV patients. Our findings implicate that Dll4 in monocyte and plasma exDll4 could be potential biomarkers of HIV-associated CVD. Targeting monocyte Dll4 could be a novel therapeutic strategy for CVD.

## Nonstandard Abbreviations and Acronyms

CVD: Cardiovascular disease
FL: Full length
LPS: Lipopolysaccharides
HIV: Human immunodeficiency virus
HUVEC: Human umbilical vein endothelial cell
LPS: Lipopolysaccharides
NICD: Notch intracellular domain
NTM: Notch transmembrane domain
PBMC: peripheral blood mononuclear cell
PLWH: People live with HIV
TNFα: Tumor necrosis factor alpha

## Supplementary figures

**Figure S1.**
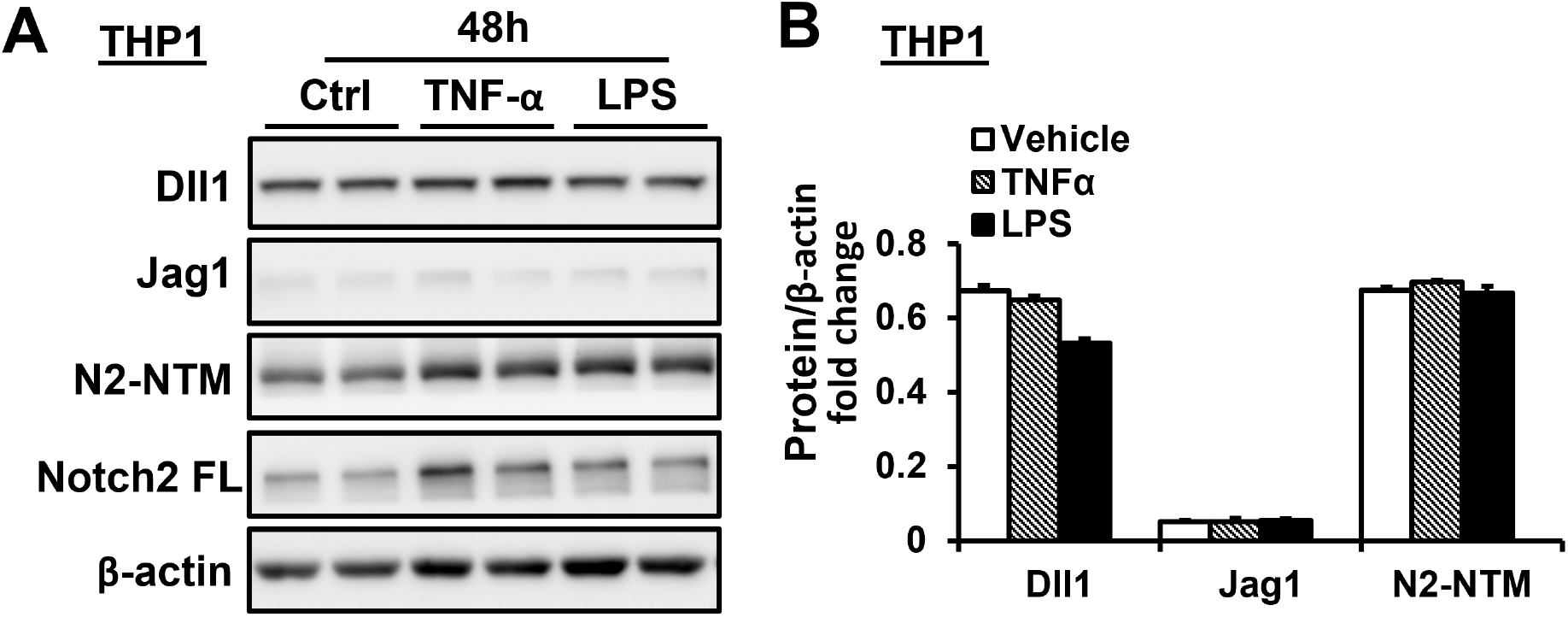
LPS and TNFα did not affect Dll1, Jag1 expression and Notch2 activation in monocytes. **(A-B)** Dll1, Jag1, N2-NTM and Notch2-full length (FL) were detected in THP1 cells with or without TNFα or LPS stimulation at 48h (n=4) by western blot **(A)** and quantification **(B)**. Proteins normalized to β-actin.

**Figure S2.**
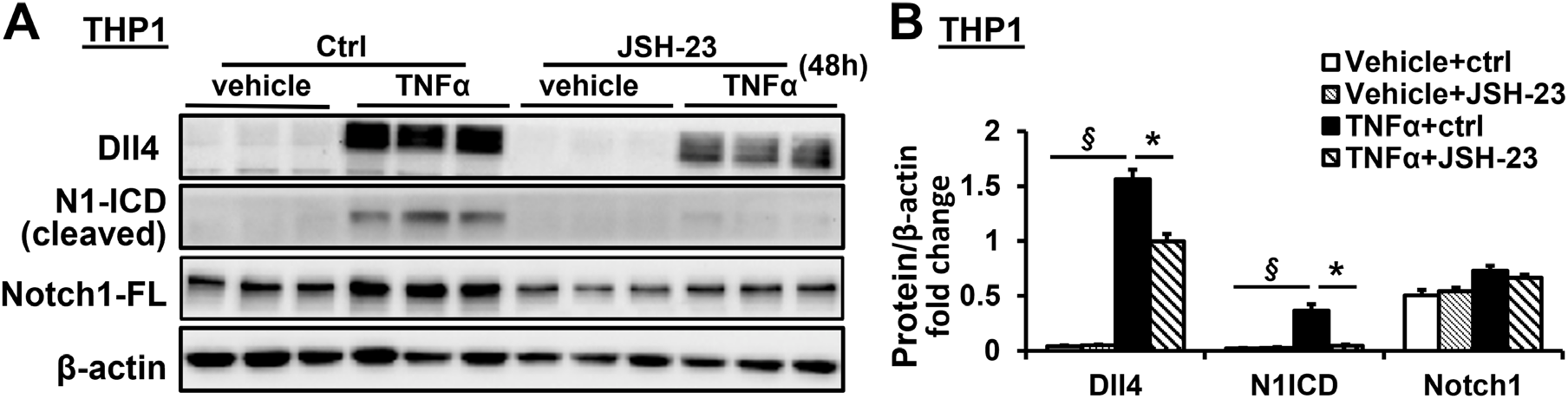
TNFα induced Dll4 expression in monocytes via NF-κB pathway. **(A-B)** Dll4 expression in monocytes exposed to proinflammatory stimuli TNFα was significantly inhibited by JSH-23 (10µM), an inhibitor of nuclear factor-κB (NF-κB), subsequently inhibited downstream Notch1 activation in THP1 cells. After inhibition of NF-κB signaling, Dll4, N1-ICD and Notch1-full length (FL) were detected in THP1 cells with or without TNFα stimulation at 48h (n=4) by western blot **(A)** and quantification **(B)**. Proteins normalized to β-actin. Data shown as Mean±SEM; *P* values were calculated using the Student’s *t*-test, and \**P*<0.05 compared with control group. § indicates significant difference between vehicle and TNFα treated control groups.

## Notes

### Competing Interest Statement

The authors have declared no competing interest.

## References

1. Shah ASV, Stelzle D, Lee KK, Beck EJ, Alam S, Clifford S, Longenecker CT, Strachan F, Bagchi S, Whiteley W, et al. Global Burden of Atherosclerotic Cardiovascular Disease in People Living With HIV: Systematic Review and Meta-Analysis. Circulation. 2018;138:1100–1112. doi: 10.1161/CIRCULATIONAHA.117.033369

2. Subramanian S, Tawakol A, Burdo TH, Abbara S, Wei J, Vijayakumar J, Corsini E, Abdelbaky A, Zanni MV, Hoffmann U, et al. Arterial inflammation in patients with HIV. JAMA. 2012;308:379–386. doi: 10.1001/jama.2012.6698

3. Tawakol A, Lo J, Zanni MV, Marmarelis E, Ihenachor EJ, MacNabb M, Wai B, Hoffmann U, Abbara S, Grinspoon S. Increased arterial inflammation relates to high-risk coronary plaque morphology in HIV-infected patients. J Acquir Immune Defic Syndr. 2014;66:164–171. doi: 10.1097/QAI.0000000000000138

4. Neuhaus J, Jacobs DR, Jr., Baker JV, Calmy A, Duprez D, La Rosa A, Kuller LH, Pett SL, Ristola M, Ross MJ, et al. Markers of inflammation, coagulation, and renal function are elevated in adults with HIV infection. J Infect Dis. 2010;201:1788–1795. doi: 10.1086/652749

5. Burdo TH, Lo J, Abbara S, Wei J, DeLelys ME, Preffer F, Rosenberg ES, Williams KC, Grinspoon S. Soluble CD163, a novel marker of activated macrophages, is elevated and associated with noncalcified coronary plaque in HIV-infected patients. J Infect Dis. 2011;204:1227–1236. doi: 10.1093/infdis/jir520

6. Kelesidis T, Kendall MA, Yang OO, Hodis HN, Currier JS. Biomarkers of microbial translocation and macrophage activation: association with progression of subclinical atherosclerosis in HIV-1 infection. J Infect Dis. 2012;206:1558–1567. doi: 10.1093/infdis/jis545

7. Sandler NG, Wand H, Roque A, Law M, Nason MC, Nixon DE, Pedersen C, Ruxrungtham K, Lewin SR, Emery S, et al. Plasma levels of soluble CD14 independently predict mortality in HIV infection. J Infect Dis. 2011;203:780–790. doi: 10.1093/infdis/jiq118

8. Liu H, Shi B, Huang CC, Eksarko P, Pope RM. Transcriptional diversity during monocyte to macrophage differentiation. Immunol Lett. 2008;117:70–80. doi: 10.1016/j.imlet.2007.12.012

9. Martinez FO, Gordon S, Locati M, Mantovani A. Transcriptional profiling of the human monocyte-to-macrophage differentiation and polarization: new molecules and patterns of gene expression. J Immunol. 2006;177:7303–7311. doi: 10.4049/jimmunol.177.10.7303

10. Dong C, Zhao G, Zhong M, Yue Y, Wu L, Xiong S. RNA sequencing and transcriptomal analysis of human monocyte to macrophage differentiation. Gene. 2013;519:279–287. doi: 10.1016/j.gene.2013.02.015

11. Takeshita K, Satoh M, Ii M, Silver M, Limbourg FP, Mukai Y, Rikitake Y, Radtke F, Gridley T, Losordo DW, et al. Critical role of endothelial Notch1 signaling in postnatal angiogenesis. Circ Res. 2007;100:70–78. doi: 10.1161/01.RES.0000254788.47304.6e

12. Nichols JT, Miyamoto A, Olsen SL, D’Souza B, Yao C, Weinmaster G. DSL ligand endocytosis physically dissociates Notch1 heterodimers before activating proteolysis can occur. Journal of Cell Biology. 2007;176:445–458. doi: 10.1083/jcb.200609014

13. Duarte A, Hirashima M, Benedito R, Trindade A, Diniz P, Bekman E, Costa L, Henrique D, Rossant J. Dosage-sensitive requirement for mouse Dll4 in artery development. Genes Dev. 2004;18:2474–2478. doi: 10.1101/gad.1239004

14. Shutter JR, Scully S, Fan W, Richards WG, Kitajewski J, Deblandre GA, Kintner CR, Stark KL. Dll4, a novel Notch ligand expressed in arterial endothelium. Genes Dev. 2000;14:1313–1318.

15. Fung E, Tang SM, Canner JP, Morishige K, Arboleda-Velasquez JF, Cardoso AA, Carlesso N, Aster JC, Aikawa M. Delta-like 4 induces notch signaling in macrophages: implications for inflammation. Circulation. 2007;115:2948–2956. doi: 10.1161/CIRCULATIONAHA.106.675462

16. Palaga T, Buranaruk C, Rengpipat S, Fauq AH, Golde TE, Kaufmann SH, Osborne BA. Notch signaling is activated by TLR stimulation and regulates macrophage functions. Eur J Immunol. 2008;38:174–183. doi: 10.1002/eji.200636999

17. Monsalve E, Ruiz-Garcia A, Baladron V, Ruiz-Hidalgo MJ, Sanchez-Solana B, Rivero S, Garcia-Ramirez JJ, Rubio A, Laborda J, Diaz-Guerra MJ. Notch1 upregulates LPS-induced macrophage activation by increasing NF-kappaB activity. Eur J Immunol. 2009;39:2556–2570. doi: 10.1002/eji.200838722

18. Fukuda D, Aikawa E, Swirski FK, Novobrantseva TI, Kotelianski V, Gorgun CZ, Chudnovskiy A, Yamazaki H, Croce K, Weissleder R, et al. Notch ligand delta-like 4 blockade attenuates atherosclerosis and metabolic disorders. Proc Natl Acad Sci U S A. 2012;109:E1868–1877. doi: 10.1073/pnas.1116889109

19. Pagie S, Gerard N, Charreau B. Notch signaling triggered via the ligand DLL4 impedes M2 macrophage differentiation and promotes their apoptosis. Cell Commun Signal. 2018;16:4. doi: 10.1186/s12964-017-0214-x

20. Fukuda D, Aikawa M. Expanding role of delta-like 4 mediated notch signaling in cardiovascular and metabolic diseases. Circ J. 2013;77:2462–2468. doi: 10.1253/circj.cj-13-0873

21. Nakano T, Fukuda D, Koga J, Aikawa M. Delta-Like Ligand 4-Notch Signaling in Macrophage Activation. Arterioscler Thromb Vasc Biol. 2016;36:2038–2047. doi: 10.1161/ATVBAHA.116.306926

22. Pagie S, Gérard N, Charreau B. Notch signaling triggered via the ligand DLL4 impedes M2 macrophage differentiation and promotes their apoptosis. Cell communication and signaling : CCS. 2018;16:4. doi: 10.1186/s12964-017-0214-x

23. Brenchley JM, Price DA, Schacker TW, Asher TE, Silvestri G, Rao S, Kazzaz Z, Bornstein E, Lambotte O, Altmann D, et al. Microbial translocation is a cause of systemic immune activation in chronic HIV infection. Nat Med. 2006;12:1365–1371. doi: 10.1038/nm1511

24. Mutlu EA, Keshavarzian A, Losurdo J, Swanson G, Siewe B, Forsyth C, French A, Demarais P, Sun Y, Koenig L, et al. A compositional look at the human gastrointestinal microbiome and immune activation parameters in HIV infected subjects. PLoS Pathog. 2014;10:e1003829. doi: 10.1371/journal.ppat.1003829

25. Nowak P, Troseid M, Avershina E, Barqasho B, Neogi U, Holm K, Hov JR, Noyan K, Vesterbacka J, Svard J, et al. Gut microbiota diversity predicts immune status in HIV-1 infection. AIDS. 2015;29:2409–2418. doi: 10.1097/QAD.0000000000000869

26. Xu W, Luo Z, Alekseyenko AV, Martin L, Wan Z, Ling B, Qin Z, Heath SL, Maas K, Cong X, et al. Distinct systemic microbiome and microbial translocation are associated with plasma level of anti-CD4 autoantibody in HIV infection. Sci Rep. 2018;8:12863. doi: 10.1038/s41598-018-31116-y

27. Murray KD, Singh MV, Zhuang Y, Uddin MN, Qiu X, Weber MT, Tivarus ME, Wang HZ, Sahin B, Zhong J, et al. Pathomechanisms of HIV-Associated Cerebral Small Vessel Disease: A Comprehensive Clinical and Neuroimaging Protocol and Analysis Pipeline. Front Neurol. 2020;11:595463. doi: 10.3389/fneur.2020.595463

28. Pang J, Yan C, Natarajan K, Cavet ME, Massett MP, Yin G, Berk BC. GIT1 mediates HDAC5 activation by angiotensin II in vascular smooth muscle cells. Arterioscler Thromb Vasc Biol. 2008;28:892–898. doi: 10.1161/ATVBAHA.107.161349

29. Pang J, Xu X, Getman MR, Shi X, Belmonte SL, Michaloski H, Mohan A, Blaxall BC, Berk BC. G protein coupled receptor kinase 2 interacting protein 1 (GIT1) is a novel regulator of mitochondrial biogenesis in heart. J Mol Cell Cardiol. 2011;51:769–776. doi: 10.1016/j.yjmcc.2011.06.020

30. Patel AA, Zhang Y, Fullerton JN, Boelen L, Rongvaux A, Maini AA, Bigley V, Flavell RA, Gilroy DW, Asquith B, et al. The fate and lifespan of human monocyte subsets in steady state and systemic inflammation. J Exp Med. 2017;214:1913–1923. doi: 10.1084/jem.20170355

31. Brown SB, Savill J. Phagocytosis triggers macrophage release of Fas ligand and induces apoptosis of bystander leukocytes. J Immunol. 1999;162:480–485.

32. Majumder S, Zhu G, Xu X, Senchanthisai S, Jiang D, Liu H, Xue C, Wang X, Coia H, Cui Z, et al. G-Protein-Coupled Receptor-2-Interacting Protein-1 Controls Stalk Cell Fate by Inhibiting Delta-like 4-Notch1 Signaling. Cell Rep. 2016;17:2532–2541. doi: 10.1016/j.celrep.2016.11.017

33. Wong KL, Tai JJ, Wong WC, Han H, Sem X, Yeap WH, Kourilsky P, Wong SC. Gene expression profiling reveals the defining features of the classical, intermediate, and nonclassical human monocyte subsets. Blood. 2011;118:e16–31. doi: 10.1182/blood-2010-12-326355

34. Yang J, Zhang L, Yu C, Yang XF, Wang H. Monocyte and macrophage differentiation: circulation inflammatory monocyte as biomarker for inflammatory diseases. Biomarker research. 2014;2:1. doi: 10.1186/2050-7771-2-1

35. Cros J, Cagnard N, Woollard K, Patey N, Zhang SY, Senechal B, Puel A, Biswas SK, Moshous D, Picard C, et al. Human CD14dim monocytes patrol and sense nucleic acids and viruses via TLR7 and TLR8 receptors. Immunity. 2010;33:375–386. doi: 10.1016/j.immuni.2010.08.012

36. Sampath P, Moideen K, Ranganathan UD, Bethunaickan R. Monocyte Subsets: Phenotypes and Function in Tuberculosis Infection. Front Immunol. 2018;9:1726. doi: 10.3389/fimmu.2018.01726

37. Ziegler-Heitbrock L. The CD14+ CD16+ blood monocytes: their role in infection and inflammation. J Leukoc Biol. 2007;81:584–592. doi: 10.1189/jlb.0806510

38. So J, Tai AK, Lichtenstein AH, Wu D, Lamon-Fava S. Sexual dimorphism of monocyte transcriptome in individuals with chronic low-grade inflammation. Biol Sex Differ. 2021;12:43. doi: 10.1186/s13293-021-00387-y

39. Bongen E, Lucian H, Khatri A, Fragiadakis GK, Bjornson ZB, Nolan GP, Utz PJ, Khatri P. Sex Differences in the Blood Transcriptome Identify Robust Changes in Immune Cell Proportions with Aging and Influenza Infection. Cell Rep. 2019;29:1961–1973 e1964. doi: 10.1016/j.celrep.2019.10.019

40. Beenakker KGM, Westendorp RGJ, de Craen AJM, Chen S, Raz Y, Ballieux B, Nelissen R, Later AFL, Huizinga TW, Slagboom PE, et al. Men Have a Stronger Monocyte-Derived Cytokine Production Response upon Stimulation with the Gram-Negative Stimulus Lipopolysaccharide than Women: A Pooled Analysis Including 15 Study Populations. J Innate Immun. 2020;12:142–153. doi: 10.1159/000499840

41. Klein SL, Flanagan KL. Sex differences in immune responses. Nature reviews Immunology. 2016;16:626–638. doi: 10.1038/nri.2016.90

42. Varghese M, Clemente J, Lerner A, Abrishami S, Islam M, Subbaiah P, Singer K. Monocyte Trafficking and Polarization Contribute to Sex Differences in Meta-Inflammation. Frontiers in Endocrinology. 2022;13. doi: 10.3389/fendo.2022.826320

43. Hatleberg CI, Ryom L, El-Sadr W, Mocroft A, Reiss P, De Wit S, Dabis F, Pradier C, d’Arminio Monforte A, Kovari H, et al. Gender differences in the use of cardiovascular interventions in HIV-positive persons; the D:A:D Study. J Int AIDS Soc. 2018;21. doi: 10.1002/jia2.25083

44. Bermick JR, Lincoln PM, Allen RM, Kunkel SL, Schaller MA. Elevated Notch ligands in serum are associated with HIV/TB coinfection. J Clin Tuberc Other Mycobact Dis. 2021;24:100258. doi: 10.1016/j.jctube.2021.100258

45. Schaller MA, Allen RM, Kimura S, Day CL, Kunkel SL. Systemic Expression of Notch Ligand Delta-Like 4 during Mycobacterial Infection Alters the T Cell Immune Response. Front Immunol. 2016;7:527. doi: 10.3389/fimmu.2016.00527

